# Music-induced analgesia in chronic pain conditions: a systematic review and meta-analysis

**DOI:** 10.1101/105148

**Authors:** Eduardo A. Garza-Villarreal, Victor Pando, Peter Vuust, Christine Parsons

## Abstract

Music is increasingly used as an adjuvant for chronic pain management as it is not invasive, inexpensive, and patients usually report positive experiences with it. However, little is known about its clinical efficacy in chronic pain patients. In this systematic review and meta-analysis, we investigated randomized controlled trials (RCTs) of adult patients that reported any type of music intervention for chronic pain, chosen by the researcher or patient, lasting for any duration. Searches were performed using PsycINFO, Scopus and PubMed for RTCs published until the end of May 2016. The primary outcome was reduction in self-reported pain using a standardized pain measurement instrument reported post-intervention. The secondary outcomes were: quality of life measures, depression and anxiety measures, among others. The study was pre-registered with PROSPERO (CRD42016039837) and the meta-analysis was done using RevMan. We identified 768 titles and abstracts, and we included 14 RTCs that fulfilled our criteria. The sample size of the studies varied between 25 and 200 participants. We found that music reduced chronic pain, and depression, with higher effect size on pain and depression. We also found music had a higher effect when the participant chose the music in contrast with researcher-chosen music. The sample size of RCTs was small and sometimes with different outcome measures. Our analysis suggests that music may be beneficial as an adjuvant for chronic pain patients, as it reduces self-reported pain and its common co-morbidities. Importantly, the analgesic effect of music appears higher with self-chosen over researcher-chosen music.

## Introduction

Chronic pain (CP) is highly prevalent worldwide, although estimates vary considerably across countries [12,14,27]. It is an important socio-economical and health problem due to the secondary disability, co-morbidities such as anxiety, depression and suicide, as well as the high rate of dependency on opioid painkillers. It is estimated that between 10-50% of patients with CP suffer mild to severe secondary disability, and CP is one of the leading causes of years lived with disability according to the Global Burden Disease study [39]. Furthermore, there is evidence that addiction to painkillers can be a route to heroin dependency [23]. For these reasons, there is a need for adjuvant therapies that help reducing pain, its comorbidities, and particularly to reduce or avoid painkiller dependency.

The most accepted hypothesis states that pain perception can be centrally modulated via the descending pain modulatory system (DPMS) by either inhibiting or facilitating nociceptive input at the brainstem and spinal cord level [36]. The DPMS can be affected by many intrinsic factors including: expectation, attention, context, sensitization, emotion, mood, chemical pathways (neurotransmitter dysfunction), and even genetics[4]. A number of studies have shown the analgesic effects of music in acute experimental pain [17,25,30,31]. The underlying mechanisms for the so-called “music-induced analgesia” (MIA) may not be specific to music, but surely related to the DPMS[35]. Music characteristics such as: high familiarity [25], few beats-per-minute [11], and self-chosen music [26], have been reported to elicit cognitive and emotional mechanisms, such as: distraction [22], pleasure [32], sense of control [4], and placebo-like effects [21], that may contribute to the analgesic effect, all of which can affect the DPMS. It is therefore possible that music provides an easily accessible and strong medium for top-down influence of the DPMS, thus reducing pain. If this hypothesis is correct, then characteristics of the “music treatment” such as music genre or delivery (listening or performing) may be less relevant for the analgesic effect, and personal preference may be more important instead.

The interest in music for the management of chronic pain during or after hospitalization is growing [1,15,18], but its use is far from routine. A growing number of studies have supported the use of music as an adjuvant [20,37,38].A recent meta-analysis by Hole et al 2015 published in Lancet highlighted the importance of listening to music as an aid for postoperatory recovery, as it reduces pain and anxiety. Nevertheless, an assessment of study quality and synthesis of findings is clearly warranted, especially in the subject of chronic pain. Providing evidence of its potential effect could encourage physicians and healthcare professionals to use it more widely with this population. In this systematic review and meta-analysis, we aim to assess the effect of music as an adjuvant for chronic pain, as well as the characteristics of the music, if any, with the better clinical response. For this we assessed evidence for the efficacy of music in reducing pain and comorbidities such as anxiety and depression in randomized controlled trials (RCTs) treating chronic pain patients. We also investigated a priori identified subgroups such as patient familiarity with the music, experimenter-chosen vs self-chosen music, and type of pain condition.

## Methods

This meta-analysis and systematic review followed procedures from the Cochrane Handbook for Systematic Reviews [19] and from the Center for Reviews and Dissemination (Centre for & Dissemination, 2014). The review protocol was registered with PROSPERO No. [CRD42016039837].

### Search Strategy

Searches were performed using PsycINFO, Scopus and PubMed for randomized controlled trials (RCTs) published until the end of May of 2016, which reported on a music intervention for chronic pain. The search terms were prevalent chronic pain conditions obtained from the IASP (http://www.iasp-pain.org/PublicationsNews/Content.aspx?ItemNumber=1673&navItemNumber=677) and MeSH terms for these (for details and search strings see Supplementary Materials). Only RCTs with adults aged between 18-70 years old were included. Exclusion criteria were: not using an RCT design, acute rather than chronic pain conditions and testing a pediatric population. At full text review, studies were checked to ensure reporting of results from unique, non-overlapping participants. We included studies that reported any type of music intervention for chronic pain: active music playing, listening to music (passive), or 'music medicine', chosen by the researcher or patient, lasting for any duration. Chronic pain was defined as pain persisting or recurring for more than 12 weeks [24]. Pain conditions were broadly defined to encompass all types of pain according to the terms by the International Association for the Study of Pain. The primary outcome was reduction in self-reported pain using standardized pain measurement instruments such as: Visual Analogue Scale, Numerical Rating Scale, The McGill Pain Questionnaire, West Haven-Yale Multidisciplinary Pain Inventory, reported post-intervention [6]. The secondary outcomes were: quality of life measures, psychological health (depression, anxiety measures), physical health (such as pain-related disability) reported post-intervention and longer-term intervention outcomes, if available. The comparison/control were: wait list control, no music control and active control, hence, there were no restrictions placed on the control.

### Data extraction and synthesis

Information was extracted from each study as follows: (1) The characteristics of the study where relevant, including: the year of publication, design, randomization, blinding, number of participants, attrition, type of outcome measures, overall treatment effects,(2) The characteristics of the intervention: the type of music, list of songs or any other detail on music included, the duration of the music intervention, measures of patient engagement and enjoyment and the qualifications/background of the individual(s) delivering the intervention, if relevant, (3) The characteristics of the participants: sex, age, dropouts, length of pain condition, co-morbid conditions. To avoid risk of bias, two reviewers (VP, EGV) independently assessed the methodological quality of the included RCTs using the Cochrane Risk of Bias (RoB) tool. We also assessed for risk of publication bias. We performed a sensitivity analysis, removing studies at high risk of bias. Any disagreements that arose between the reviewers was resolved through discussion, or with a third reviewer (CEP).

To analyze the association between intervention and outcome, and to avoid variability from the different pain measurement instruments and missing data, we used the primary outcome post-intervention as reported by the study investigators and calculated the effect size. Only in one study [34], the post-intervention improvement was shown as an increase in the pain variable, hence the effect size was calculated backwards. If studies showed pain reduction as a secondary outcome of interest, we considered it the primary outcome for our analyses. Statistical analyses, including pooled mean home practice data and meta regression and risk of bias, was conducted using the software RevMan 5.3 [10]. An estimation of heterogeneity was calculated.

### Subgroup Analyses

We investigated several *a priori* identified subgroups of interest: “primary vs secondary pain”, “central vs peripheral pain”, “long-term chronic pain conditions (> 5 years) vs short-term pain conditions”, and “participant-selected vs researcher-selected music”. We conducted the subgroup analyses using difference of means. As *post-hoc*, we removed the “primary vs secondary pain” contrast because the terms are not commonly used, and we also removed “long-term vs short-term pain” contrast due to the low sample size for the later (n = 1). Instead, we included the contrast “etiology of pain” that includes IASP categories: neoplasm, degenerative/mechanical, inflammatory, unknown etiology and combined etiology (studies that included two or more pathologies). We also studied the contrast intervention “familiar vs unfamiliar music”. Finally, we added “music delivery” (recorded music, live music, or active music) as a subgroup contrast of interest.

### Study characteristics

Figure 1 shows the PRISMA flow chart for the included studies. A total of 65 studies were identified that reported a music intervention for chronic pain (N=768). Of those, 14 studies fulfilled our criteria and/or provided data when contacted. 11 studies investigated pain reduction as a primary outcome, while 3 investigated pain as a secondary outcome. The characteristics of the included studies are shown in Table 1. The sample size of the studies varied between 25 and 200 participants (M = 84; SD = 47). The total sample of participants included for this review and meta-analysis was n = 1,178. The majority of included studies examined patients experiencing either cancer pain (4/14) or fibromyalgia (3/14), but a variety of other patient groups were also examined (palliative care, osteoarthritis, multiple sclerosis, chronic non-malignant pain, and inflammatory bowel disease). The mean age of included participants was 55 years (± 10.8), and the mean length of pain condition was 7.3 years (± 4.1). The mean music duration in the included studies was 30 minutes (± 10.05), 80% of them between 20 - 30 minutes. The majority of studies with recorded music intervention used headphones for delivery (7/11), while the others used CD players.

**Table 1.** Characteristics of studies included in the meta-analysis evaluating music for chronic pain.

| Study | Country | Participants | Sample | Intervention | Control | Qualification | Tools |
| --- | --- | --- | --- | --- | --- | --- | --- |
| Alparslan et al. (2016) | Turkey | Fibromyalgia | N=37, mean age=43.59, 95% female | Recorded Music | No Music | Participant-administered | pVAS (0-10) |
| Burrai et al. (2014). | Italy | Cancer | N=52, mean age=64.5, 83% female | Live Music | No Music, Rest | Trained nurse | mVAS (0-10), pVAS (0-10, BP) |
| Clark et al. (2006). | USA | Cancer | N=63, mean age=57.79, 38% female | Recorded Music | No Music | Music therapist | dNRS (0-10), pNRS (0-10), HADS, POMS |
| Grape et al. (2009). | Sweden | Irritable Bowel Syndrome | N=55, mean age=not reported, sex not reported | Active Music | No Choir Singing | Participant-administered | VAS (0-10), BP |
| Guétin et al. (2012). | France | CNMP | N=87, mean age=48.85, 78% female | Recorded Music | No Music, Standard Care | Trained nurse | VAS (0-10), HADS, pNRS (0-10) |
| Gutgsell et al. (2013). | USA | Palliative care | N=200, mean age=56.09, 69% female | Live Music | No Music, Rest | Music therapist | pNRS (0-10), FLACCs, FPS (0-5) |
| Horne-Thompson (2008). | Australia | Palliative care | N=25, mean age=73.9, 44% female | Recorded Music | Reading, conversation | Music therapist | ESAS (0-10), BP |
| Huang et al. (2010). | Taiwan | Cancer | N=126, mean age=54, 29% female | Recorded Music | No Music, Rest | Trained nurse | VAS (0-100) |
| Li et al. (2011). | China | Cancer | N=120, mean age=45.01, 100% female | Recorded Music | No Music | Trained researcher | VAS (0-10), PPI (0-5), PRI |
| McCaffrey (2003). | USA | Osteoarthritis | N= 66, mean age=76.58, 67% female | Recorded Music | No Music, Rest | Participant-administered | VAS (0-100), PRI |
| Onieva-Zafra et al. (2013). | Spain | Fibromyalgia | N=55, mean age=51.6, 96% female | Recorded Music | No Music | Participant-administered | VAS (0-10), McGPQ, BDI |
| Seebacher et al. (2016). | UK | Multiple Sclerosis | N=112, mean age=44.1, 76% female | Recorded Music | Metronome, No Music | Participant-administered | VAS (0-100), BP |
| Siedliecki (2006). | USA | CNMP | N=60, mean age=49.7, 77% female | Recorded Music | No Music, Standard Care | Participant-administered | VAS (0=10), McGPQ, CESD. |
| Weber et al. (2015). | Brazil | Fibromyalgia | N=120, mean age=48.27, 100% female | Recorded Music | No Music, Vibration Points | Participant-administered | FIQ, HAQ |

**Figure 1.**
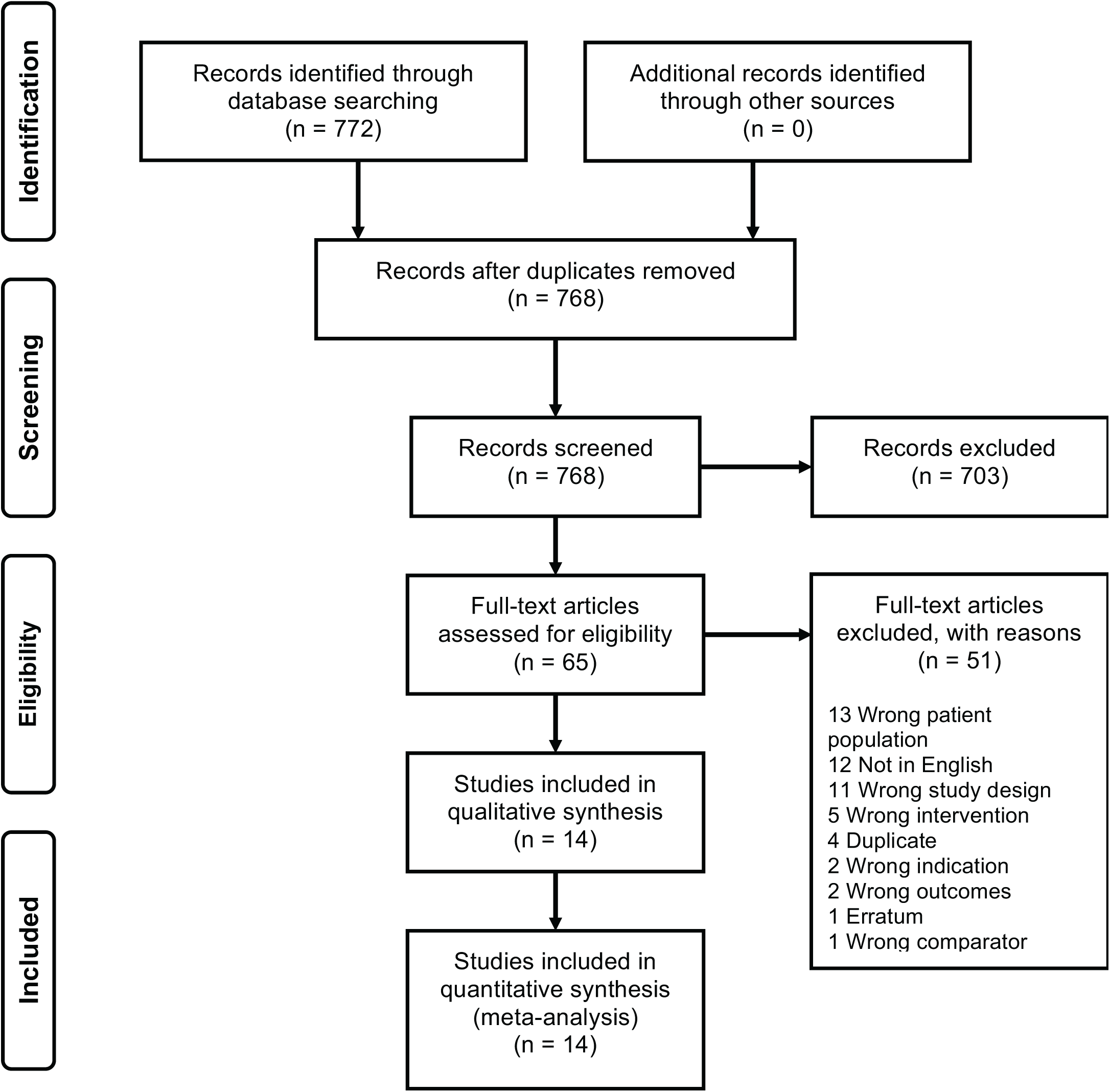
Flow diagram for review of studies of music interventions for chronic pain.

The music delivery was performed by the participant in most studies (7/14), while in others the intervention was delivered by a trained researcher, a trained nurse, or a music therapist. Music was delivered via recorded tapes in the majority of cases (11/14), or live music (2/14). One study was considered “active music”, as choir singing was the intervention (1/14). Only 2 studies (14%) reported “music enjoyment” as a variable, both reporting patients’ liking of music as > 90% (cite). Neither study stated the method used to assess music enjoyment. Timing of delivery was anytime of the day, including moments with increased perceived pain. We identified some genres of music delivered: Chinese classical music, Swedish songs, Taiwanese folk songs, buddhist music, classical western music, jazz, pop, as well as instrumental and ambient music such as: ocean drum, harp, piano, orchestra, water and wave sounds. Therefore, music genre varied greatly between studies.

Comparator/control descriptions varied and include different ones: no music (11/14), tactile touch (1/14), conversation (1/14), and routine patient care (1/14). Pain was usually measured with VAS (10/11), but other scales were used such as numerical rating scales (NRS) (1/14), functional pain scale (FPS) (1/14), and pain rating index (PRI) (1/14). Secondary outcomes were measured with a variety of scales, such as: the Hospital Anxiety and Depression Scale (HADS) [40], the Edmonton Symptom Assessment System (ESAS) [28], the Beck Depression Inventory (BDI) [2], the Fibromyalgia Impact Questionnaire (FIQ) [3], and the Health Assessment Questionnaire (HAQ) [5].

## Results

We identified a total of 768 titles and abstracts, of which we reviewed 65 studies at full text that reported music intervention for chronic pain (see PRISMA Figure 1). We included 14 RCTs in the final qualitative and quantitative synthesis. Of the 14 studies included in the review, 11 investigated pain reduction as a primary outcome, while 3 investigated pain reduction as a secondary outcome. Other secondary outcomes included: anxiety, depression, fatigue, quality of life, disability and biological parameters. We found that music reduced chronic pain in general (14 RCTs, SMD −0.60 [ −0.72, - 0.48], Z = 9.81, p < 0.001) (Figure 2), anxiety (4 RCTs, SMD −0.55 [ −0.80, −0.30], Z = 4.31, p < 0.001) (Figure 3), and depression (4 RCTs, SMD −0.82 [−1.08, −0.56], Z = 6.12, p < 0.001) (Figure 4). Effect size was moderate for chronic pain and anxiety, while depression had a high size effect. Heterogeneity was high for pain, anxiety, and depression, with an I^2^ = 60%, 85%, and 88% respectively. A domain-based evaluation was used as the tool for assessing risk of bias. The quality of studies was adequate for most of the included authors, with a low risk of bias overall (Figure 5). Selection bias, attrition bias, and reporting bias had a low risk given that randomization was well performed, outcome data was complete, attrition was low, and all measured outcomes were reported. We found that performance risk of bias was moderate, due to lack of blinding for 8 of the 14 studies. Assessing for selection bias, we found moderate risk because the allocation concealment was not stated (3/14), or was not done (2/14) (Figure 6).

**Figure 2.**
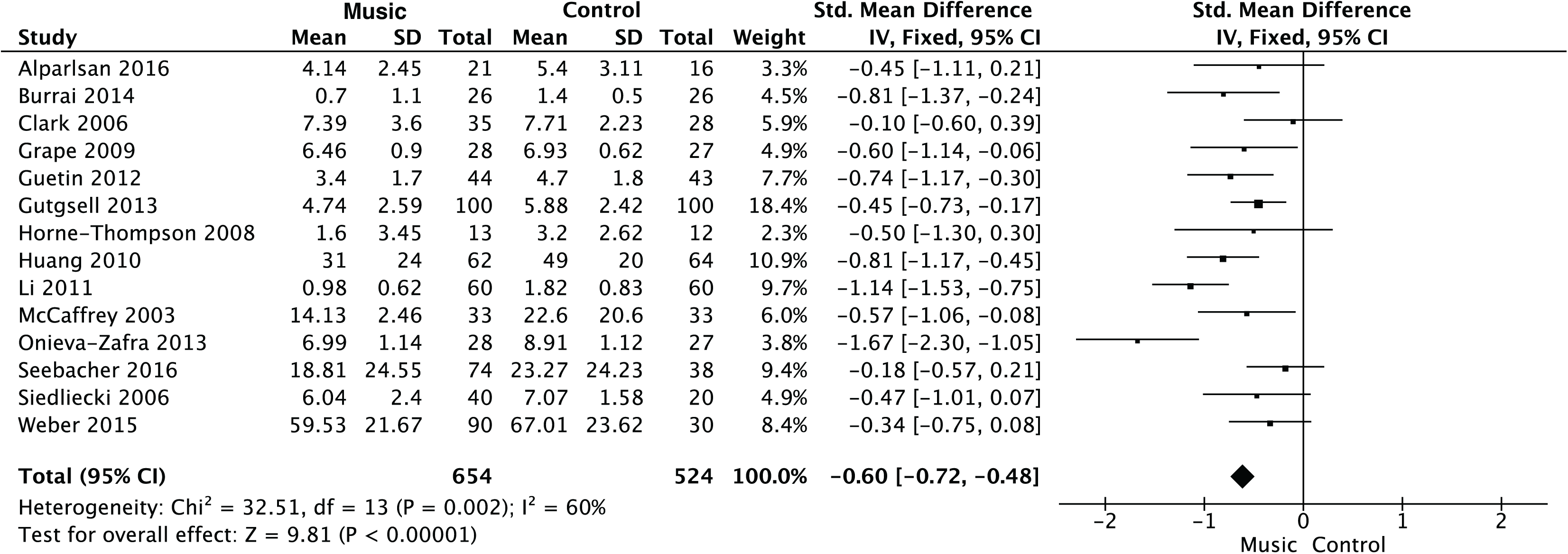
Music effects on chronic pain across all studies. SD = standard deviation, CI =
Confidence Interval, IV = inverse variance, I^2^ = Inconsistency.

**Figure 3.**
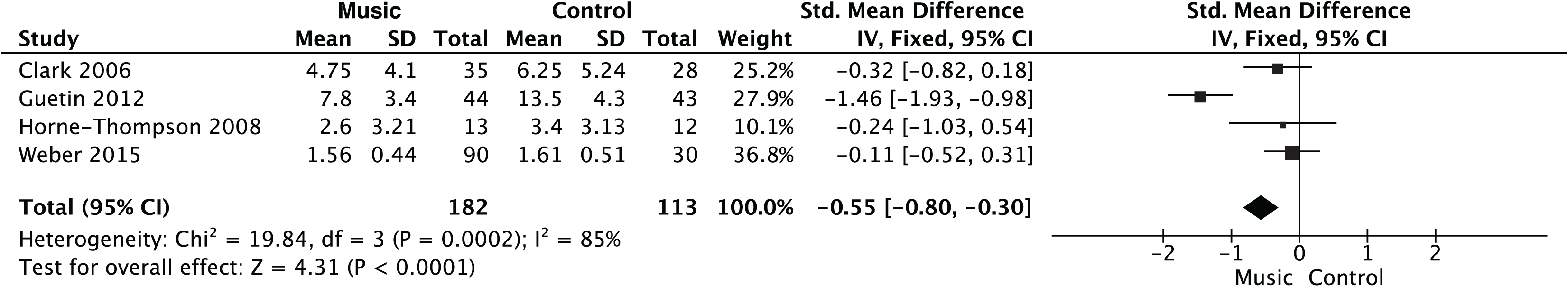
Music effects on anxiety across all studies. SD = standard deviation, CI =
Confidence Interval, IV = inverse variance, I^2^= Inconsistency.

**Figure 4.**
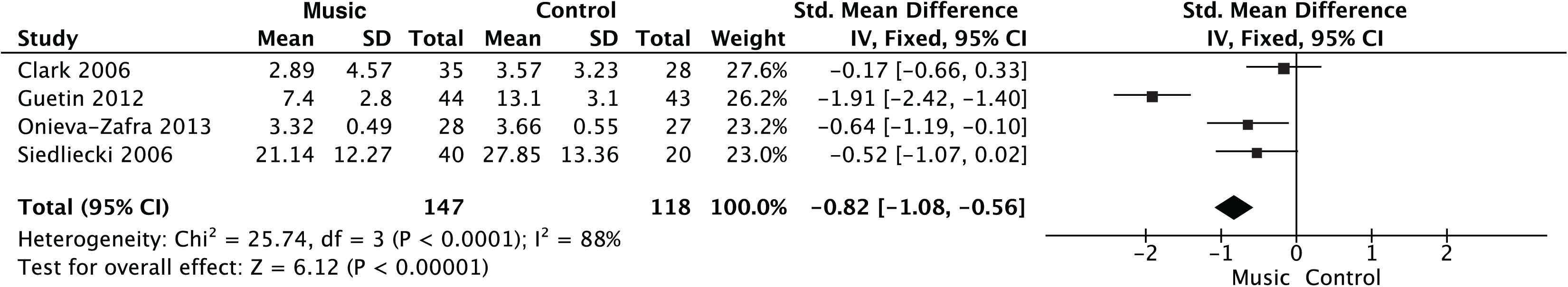
Music effects on depression across all studies. SD = standard deviation, CI =
Confidence Interval, IV = inverse variance, I^2^ = Inconsistency.

**Figure 5.**
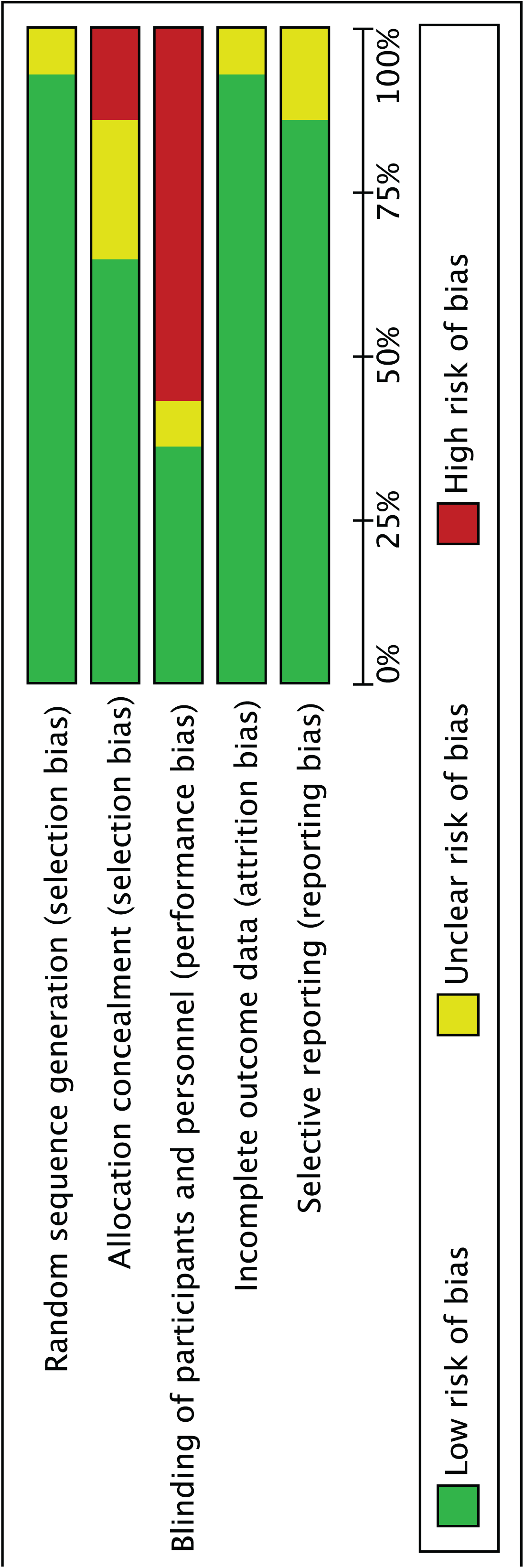
Risk of Bias across studies.

**Figure 6.**
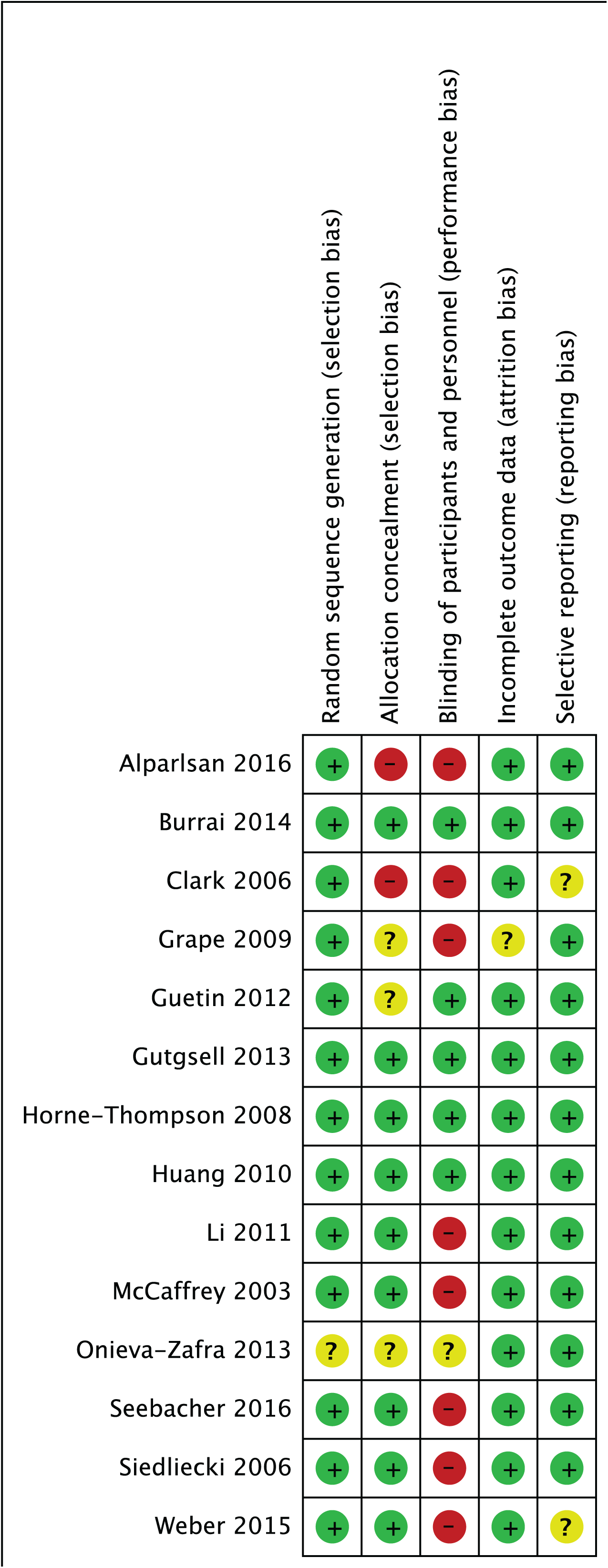
Risk of Bias in individual studies.

### Subgroup analysis

The subgroup analysis by “etiology of pain” (Figure 7) showed a non-significant (p = 0.17) higher effect size of music intervention when the pain etiology was neoplasm (4 RCTs, SMD −0.78 [-1.00, −0.56]) rather than degenerative/mechanical (2 RCTs, SMD - 0.33 [ −0.64, −0.03], inflammatory (1 RCT, SMD −0.60 [-1.14, −0.06], unknown etiology (3 RCT, SMD −0.68 [ −0.99, −0.38]), and combined etiology (4 RCT, SMD −0.52 [ −0.73, - 0.32]). The subgroup analysis “central vs peripheral pain” showed no differences of music intervention (p = 0.81) between central pain (4 RCTs, SMD −0.66 [ −0.93, −0.40]), peripheral pain (5 RCTs, SMD −0.61 [ −0.80, −0.43]), and studies with both central and peripheral pain (5 RCTs, SMD −0.56 [ −0.75, −0.36]) (Figure 8). In the “familiar vs unfamiliar music” contrast, music had a non-significant (p = 0.11) higher effect size when musical pieces were familiar (6 RCTs, SMD −0.72 [ −0.91, −0.53]), when compared to unfamiliar music (8 RCTs, SMD −0.52 [ −0.68, −0.36]) (Figure 9). In the contrast “participant-selected vs researcher-selected music”, we found a significant greater effect size (p = 0.02) when the participants chose the music (5 RCTs, SMD −0.81 [-1.02, - 0.59]) than when the researchers chose the music (9 RCTs, SMD −0.51 [ −0.65, −0.36]) (Figure 10). Examining the funnel plot (Figure 11) suggests that publication bias was not substantial as studies are evenly distributed either side of the SMD for chronic pain. Our risk of bias analysis shows low risk overall (Fig 5 & 6).

**Figure 7.**
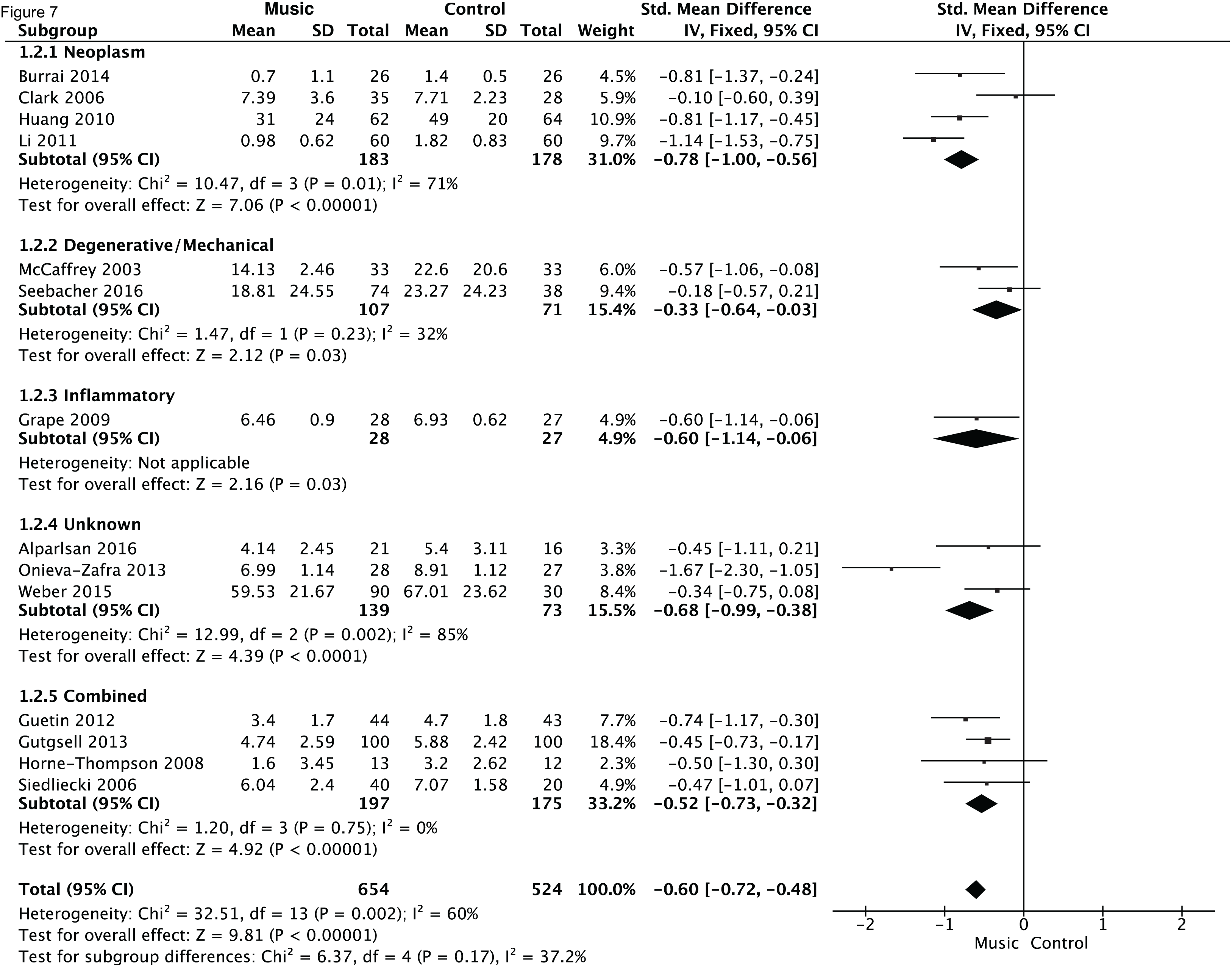
Subgroup analysis of chronic pain by Etiology of Pain. SD = standard
deviation, CI = Confidence Interval, IV = inverse variance, I^2^ = Inconsistency.

**Figure 8.**
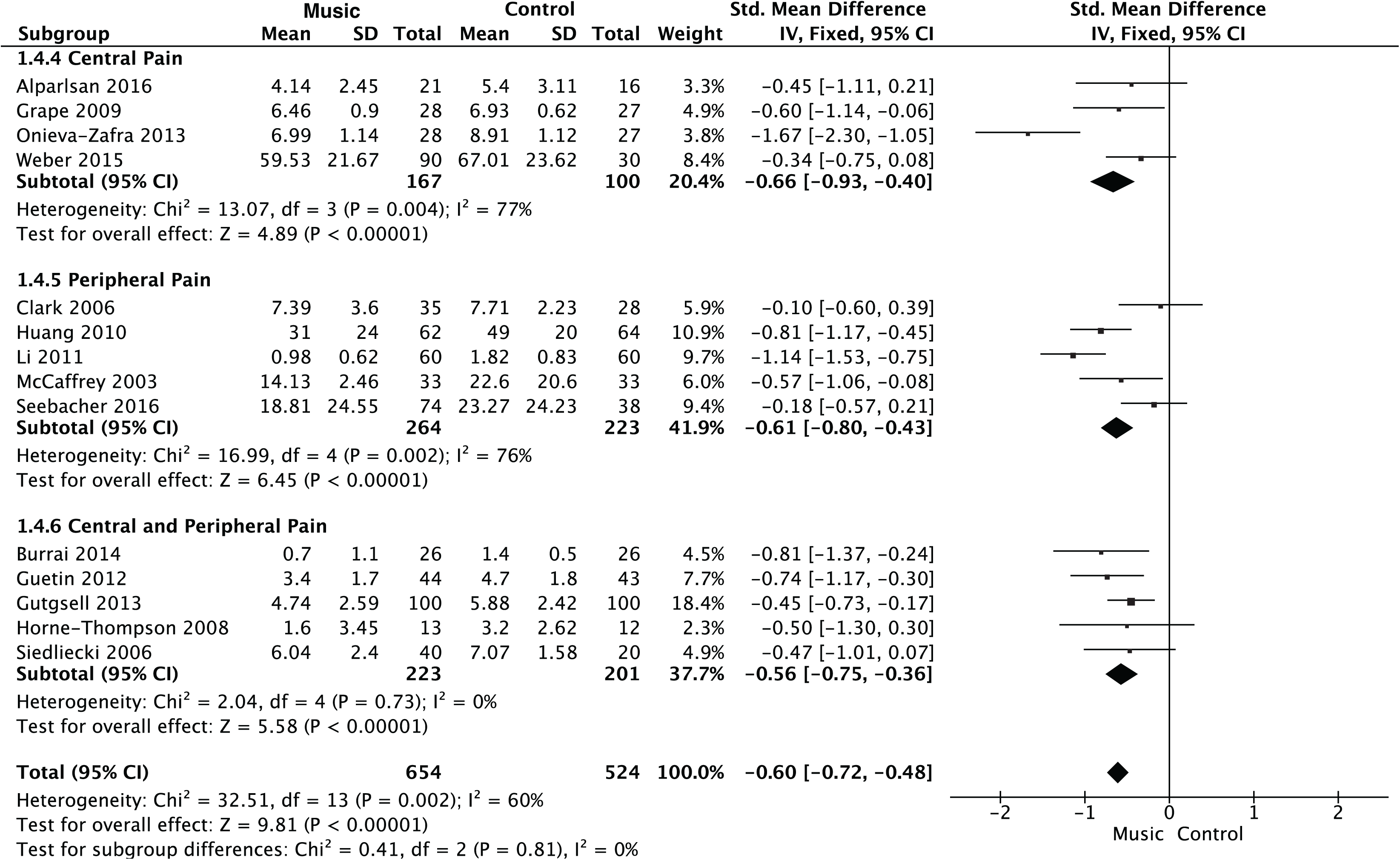
Subgroup analysis of chronic pain by Pain Location. SD = standard deviation,
CI = Confidence Interval, IV = inverse variance, I^2^ = Inconsistency.

**Figure 9.**
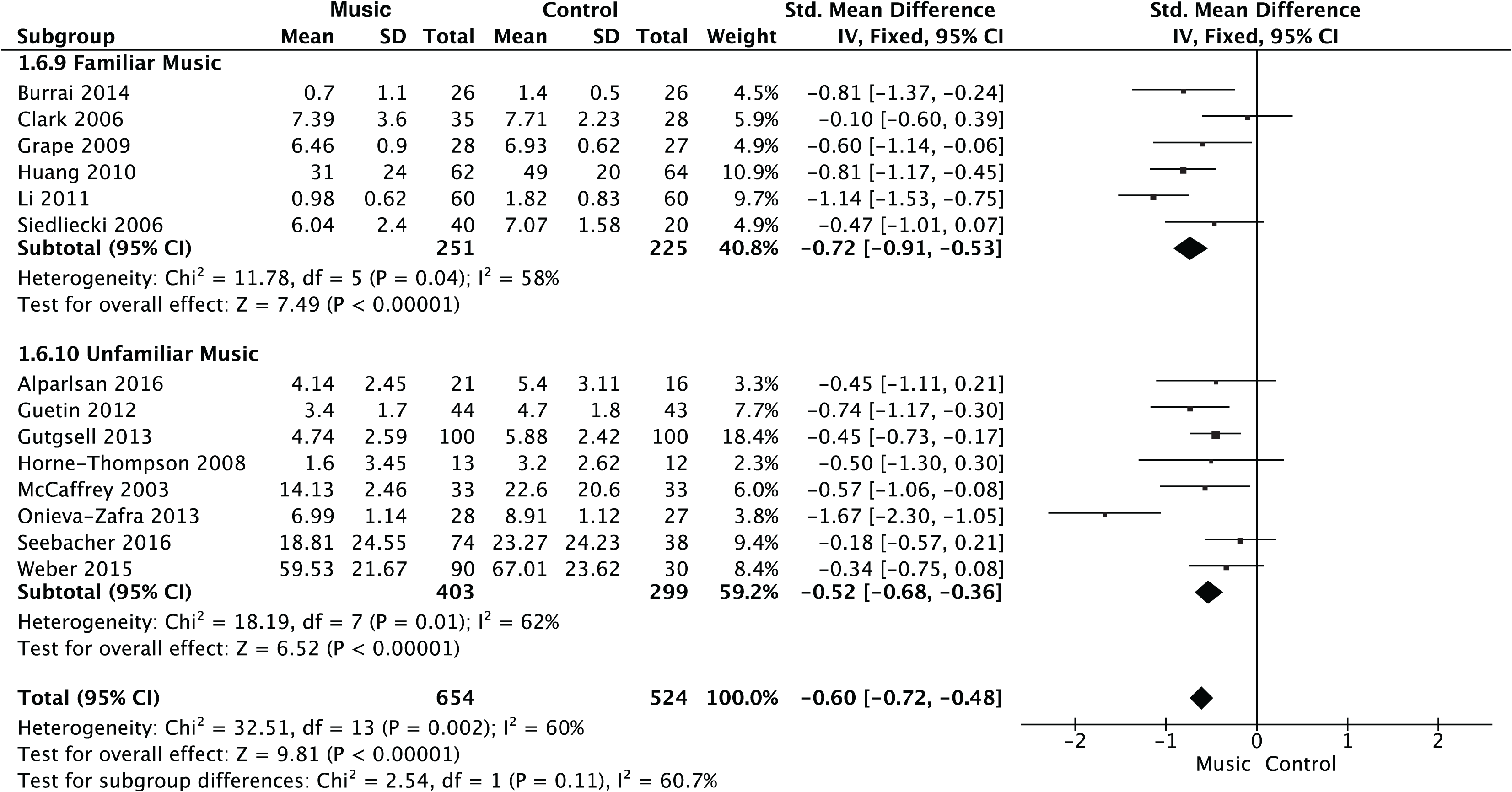
Subgroup analysis of chronic pain by Music Familiarity. SD = standard
deviation, CI = Confidence Interval, IV = inverse variance, I^2^ = Inconsistency.

**Figure 10.**
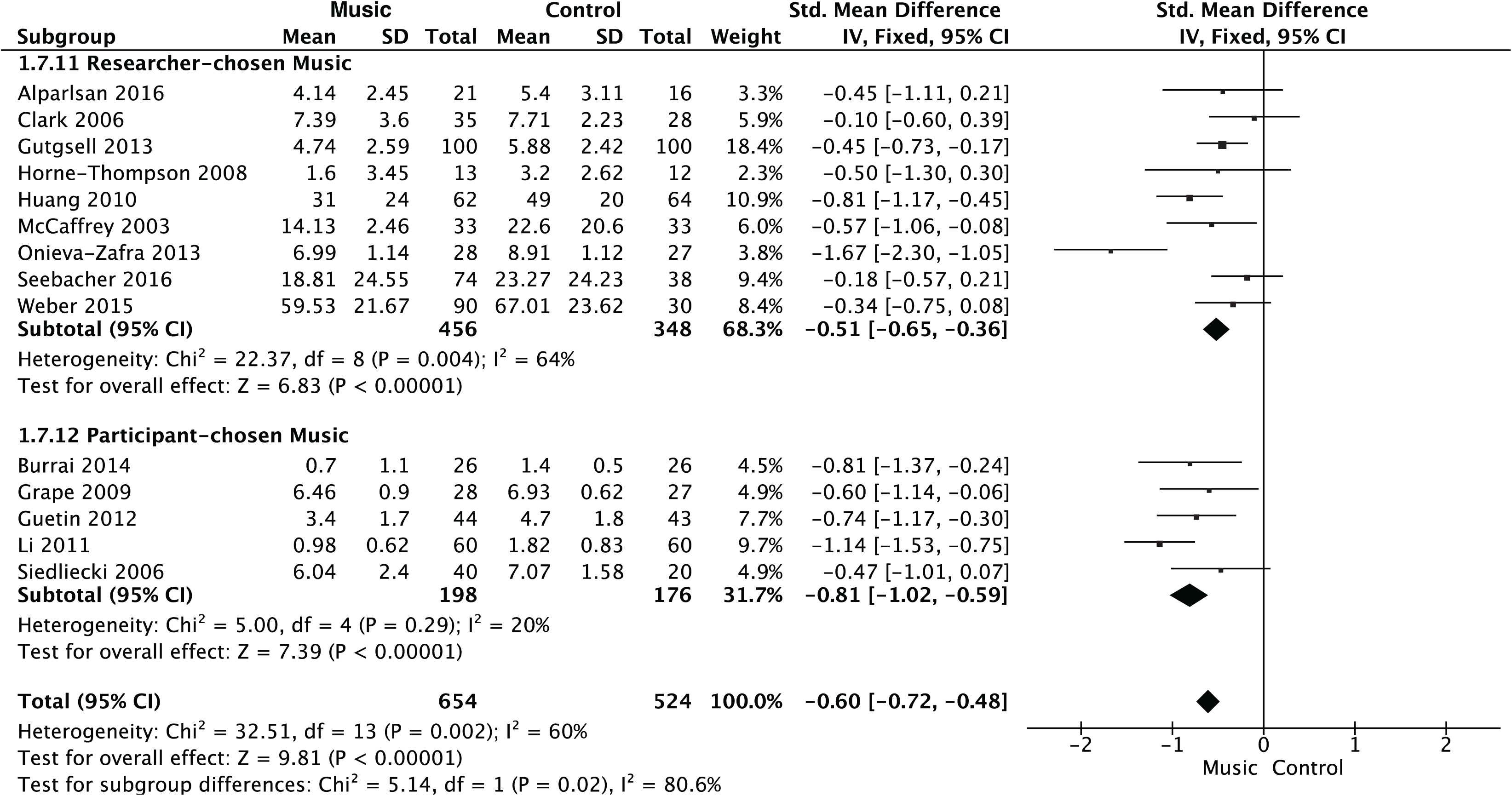
Subgroup analysis of chronic pain by Music Selection. SD = standard
deviation, CI = Confidence Interval, IV = inverse variance, I^2^ = Inconsistency.

**Figure 11.**
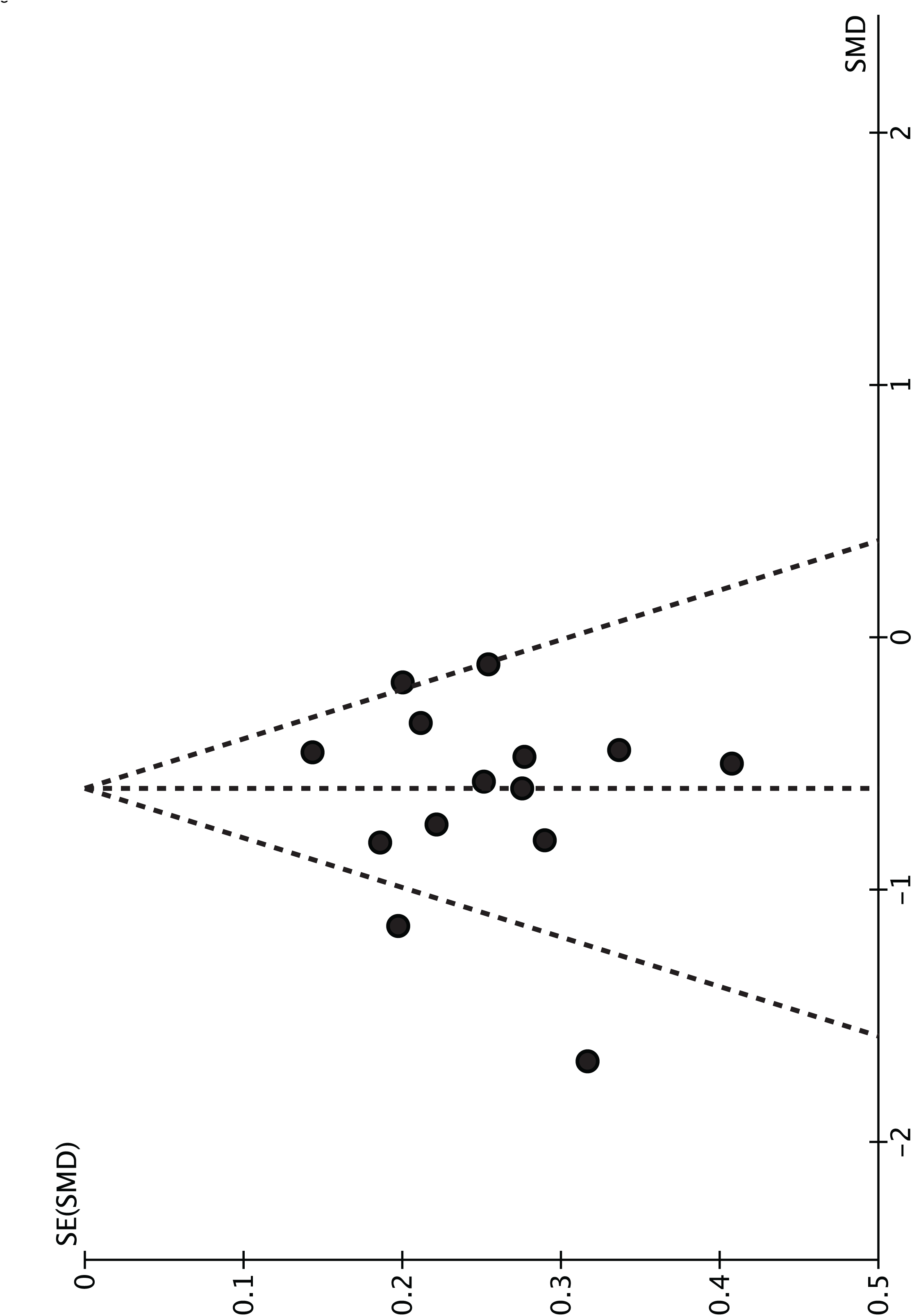
Funnel plot. SE = Standard Deviation, SMD = Standardized Mean Difference.

### Heterogeneity

Heterogeneity of effects was high across studies, probably due to several reasons: the use of different scales to measure pain, anxiety and depression; the variability in the sample sizes; the variability of the pain etiology; the variability in the duration of the study; the type of music intervention of each study; the type of music delivery (passive vs active, participant– vs researcher-chosen). In the subgroup analysis “etiology of pain” (Fig 5), the heterogeneity of studies that included combined etiology was null (Chi^2^=1.20, df=3, I^2^=0%), and in studies with degenerative/mechanical processes it was low (Chi^2^=1.47, df=1, I^2^=32%), when compared with neoplasm (Chi^2^=10.47, df=3, I^2^=71%) and unknown (Chi^2^=12.99, df=2, I^2^=85%). Studies in the combined etiology subgroup assessed pain with 0-10 VAS only, studied either CNMP (2/4) or palliative care (2/4), and evaluated both central and peripheral pain. A similar pattern occurs while analyzing heterogeneity in the subgroup “central vs peripheral pain”. Studies that included both central and peripheral pain, showed no heterogeneity (Chi^2^=2.04, df=4, I^2^=0%). These studies used only 0-10 VAS. Heterogeneity was high for studies that included central pain (Chi^2^=13.07, df=3, I^2^=77%) and peripheral pain (Chi^2^=16.99, df=4, I^2^=76%), independently.

The subgroup analysis of “familiar vs unfamiliar music” showed substantial heterogeneity in both familiar (Chi^2^=11.78, df=5, I^2^=58%), and unfamiliar (Chi^2^=18.19, df=7, I^2^=62%) music. A different pattern was shown in the subgroup “participant-chosen vs researcher-chosen music”, where heterogeneity was low for participant-chosen music (Chi^2^=5.00, df=4, I^2^=20%), compared with researcher-chosen music (Chi^2^=22.37, df=8, I^2^=64%).

## Discussion

This systematic review suggests that listening to music reduces self-reported pain, anxiety and depression symptoms in a diverse range of chronic pain patients. We also found that music helped to reduce symptoms of anxiety and depression, common disabling comorbidities in chronic pain. In a subgroup analysis, we found that the analgesic effects of music were greater for patient-chosen than researcher-chosen music.

All of the included studies reported analgesic effects of music, and in general, these effects were significant, supporting the hypothesis that music is beneficial in chronic pain conditions. This suggests that the analgesic effect of music may initiate in the brain and elicits a top-down regulation mechanism via the DPMS, as it has been suggested in fibromyalgia [17]. Only 4 studies did not report the duration of their whole intervention. The mean duration of the studies was 5 weeks (± 3), and only 1 study lasted for a whole year [9]. The duration of the analgesic effect was not directly discussed in any study, but according to their results we suggest that a daily session of 20 – 30 minutes, while experiencing pain exacerbation or not, is effective and recommended.

Reductions in anxiety and depression symptoms, aside from the pain, is an important effect, given that suicide is not uncommon. Music might improve patient coping with their condition, which at the same time may reduce the feeling of helplessness and suicidal thoughts [8]. It is not clear whether the reduction of anxiety and depression symptoms themselves could be secondary to the analgesic mechanism, but it is important to note that music seems to be positively contributing to more than one dimension and symptom of chronic pain.

### Subgroup analysis

We found no significant differences in response to music based on the etiology of pain or location of pain. Nevertheless, the studies we found did not include all types of pain (i.e. migraine) and more research is necessary to confirm our findings. One question of clear interest is how best to deliver a music intervention. In most of our included studies, the patients listened to music, and only one study used active singing [9] and two others used live music [7,18]. We therefore cannot make clear recommendations about the ideal mode of delivery of a music intervention. More studies are needed to assess if listening to recorded music (passive) provides similar benefits performing music (active) (Doelling & Poeppel 2015). We found no significant difference in pain response to familiar (6/14) and unfamiliar music studies (8/14). This result contrasts with several experimental studies showing a higher analgesic effect of familiarity [25,29]. Listening to familiar music may induce a feeling of “control” of the situation and the expectation of musical “peaks” could induce pleasure and secondary analgesia and relaxation, as well as release dopamine and endogenous opioids [33]. In an experimental study by our group we found that unfamiliar music provided less of an analgesic effect than an active math distraction [16]. However, in this synthesis, we found that the effects of music familiarity were non-significant suggesting no effect or very low effect. This may be due to the small number of studies in our sample and should be studied further.

Evidence from other studies suggests that musical genre is not important for the analgesic effect. It was not possible to study genre, however, the included studies used a somewhat wide range of music genres. In our study, self-chosen music (5/14) had a significantly higher analgesic effect than researcher-chosen music (9/14) in our study, and was in fact the only statistically significant subgroup analysis. This has been reported by several experimental studies [11] and was also shown in another meta-analysis of music in cancer patients [37]. This effect may be related to familiarity, feeling of control and pleasure, thus contributing to the analgesic effect. The use of self-chosen music may present a challenge for standardizing music treatment, and having a pool of musical choices could reduce this problem.

## Limitations

Due to the small number of studies, subgroup comparisons should be interpreted cautiously. In terms of study size, there was only one study with an n < 35, which suggests that studies overall were reasonably powered to find a moderate effect size. However, most outcome variables, primary and secondary, were measured using different instruments, which may partly explain the heterogeneity between studies. Several studies lacked complete information (i.e. pain measures and the type of music used). Also, there were many variable factors such as: the person who chose the music (participant– vs. researcher-chosen), the length of the music intervention (1 week – 12 months), the type of delivery of the music (active, passive listening, live music) and the type control condition, among others.

In our sample, no study reported adverse or negative effects with music, and this could be either because there were none, or because of inadequate reporting, which is typical for many psychological interventions [13]. Nevertheless, we found an sample of RCTs with adequate sample sizes, and overall effect sizes that provide consistent evidence in support of the reduction of pain, depression and anxiety in chronic pain conditions, hence, supporting the use of music as an adjuvant in pain medicine.

Future RCTs should aim to use active control conditions, and use more than one standard instrument (VAS, numeric, etc), and to use self-chosen music to try and avoid heterogeneity. We still do not know the precise duration of the analgesic effect of music and the dosage of music intervention to produce a positive outcome, and these may explain part of the heterogeneity. Future studies should also record medication intake to assess for any reduction in the amount of painkillers, anxiety and depression medication after music intervention. While medication information was lacking in our study, it would be of clinical interest to examine the potential reduction in pain medication intake after music intervention. Reducing the intake amount of pain medication, would improve the patients’ quality of life by avoiding secondary effects such as gastrointestinal problems and prescribed drug dependency.

## Conclusions

In this systematic review and meta-analysis, we show that music reduces pain in chronic pain conditions, and anxiety and depression symptoms, and that self-chosen music have higher analgesic effect. We suggest that music can be used as an easily administered, effective adjuvant for chronic pain and its common comorbidities. Our systematic review and meta-analysis is the most complete to date on music and chronic pain patients, given the comprehensive search terms and study choice (RCTs). More studies are necessary to untangle specific questions about the mechanisms underlying the effect of music and further studies should focus more on indirect measures such as the amount of medication taken after the music intervention.

## Acknowledgements

We would like to thank Nadia Augusto for help on the initial searches. The authors declare no conflict of interest. Christine Parsons received funding from TrygFonden. The Center for Music in the Brain at Aarhus University is funded by the Danish National Research Foundation (DNRF 117) to Peter Vuust.

## Supplementary Material: Search

### Search Strings

Pubmed results: 179

(((((((“music”[MeSH Terms] OR “music”[All Fields]) OR (“music therapy”[MeSH Terms] OR (“music”[All Fields] AND “therapy”[All Fields]) OR “music therapy”[All Fields])))) AND ((((“chronic pain”[MeSH Terms] OR (“chronic”[All Fields] AND “pain”[All Fields]) OR “chronic pain”[All Fields]) OR (“neuralgia”[MeSH Terms] OR “neuralgia”[All Fields] OR (“neuropathic”[All Fields] AND “pain”[All Fields]) OR “neuropathic pain”[All Fields]) OR (“peripheral nervous system diseases”[MeSH Terms] OR (“peripheral”[All Fields] AND “nervous”[All Fields] AND “system”[All Fields] AND “diseases”[All Fields]) OR “peripheral nervous system diseases”[All Fields] OR (“peripheral”[All Fields] AND “neuropathy”[All Fields]) OR “peripheral neuropathy”[All Fields]) OR (“neuralgia”[MeSH Terms] OR “neuralgia”[All Fields]) OR (central[All Fields] AND (“pain”[MeSH Terms] OR “pain”[All Fields])) OR (“back pain”[MeSH Terms] OR (“back”[All Fields] AND “pain”[All Fields]) OR “back pain”[All Fields]) OR (“irritable bowel syndrome”[MeSH Terms] OR (“irritable”[All Fields] AND “bowel”[All Fields] AND “syndrome”[All Fields]) OR “irritable bowel syndrome”[All Fields] OR (“irritable”[All Fields] AND “bowel”[All Fields]) OR “irritable bowel”[All Fields]) OR (inflammatory[All Fields] AND (“intestines”[MeSH Terms] OR “intestines”[All Fields] OR “bowel”[All Fields])) OR (chronic[All Fields] AND (“pelvic pain”[MeSH Terms] OR (“pelvic”[All Fields] AND “pain”[All Fields]) OR “pelvic pain”[All Fields])) OR ((“cardiovascular system”[MeSH Terms] OR (“cardiovascular”[All Fields] AND “system”[All Fields]) OR “cardiovascular system”[All Fields] OR “cardiovascular”[All Fields]) AND (“pain”[MeSH Terms] OR “pain”[All Fields])) OR ((“neoplasms”[MeSH Terms] OR “neoplasms”[All Fields] OR “cancer”[All Fields])) OR ((“tumour”[All Fields] OR “neoplasms”[MeSH Terms] OR “neoplasms”[All Fields] OR “tumor”[All Fields]) AND (“pain”[MeSH Terms] OR “pain”[All Fields])) OR ((“tumour”[All Fields] OR “neoplasms”[MeSH Terms] OR “neoplasms”[All Fields] OR “tumor”[All Fields]) AND (“pain”[MeSH Terms] OR “pain”[All Fields])) OR (“phantom limb”[MeSH Terms] OR (“phantom”[All Fields] AND “limb”[All Fields]) OR “phantom limb”[All Fields]) OR (complex[All Fields] AND regional[All Fields] AND (“pain”[MeSH Terms] OR “pain”[All Fields])) OR (“arthritis”[MeSH Terms] OR “arthritis”[All Fields]) OR (“lupus erythematosus, systemic”[MeSH Terms] OR (“lupus”[All Fields] AND “erythematosus”[All Fields] AND “systemic”[All Fields]) OR “systemic lupus erythematosus”[All Fields] OR (“systemic”[All Fields] AND “lupus”[All Fields] AND “erythematosus”[All Fields])) OR (“multiple sclerosis”[MeSH Terms] OR (“multiple”[All Fields] AND “sclerosis”[All Fields]) OR “multiple sclerosis”[All Fields]) OR (“fibromyalgia”[MeSH Terms] OR “fibromyalgia”[All Fields]) OR (“spondylitis, ankylosing”[MeSH Terms] OR (“spondylitis”[All Fields] AND “ankylosing”[All Fields]) OR “ankylosing spondylitis”[All Fields] OR (“ankylosing”[All Fields] AND “spondylitis”[All Fields])) OR (“headache”[MeSH Terms] OR “headache”[All Fields]) OR (“palliative care”[Mesh Terms] OR (“palliative”[All Fields] AND “care”[All Fields]) OR “palliative care”[All Fields]) OR (“carpal tunnel syndrome”[MeSH Terms] OR (“carpal”[All Fields] AND “tunnel”[All Fields] AND “syndrome”[All Fields]) OR “carpal tunnel syndrome”[All Fields]) OR (“glaucoma”[MeSH Terms] OR“glaucoma”[All Fields]))))))) AND ((random* AND control* AND trial) OR RCT OR clinical trial OR random alloca*)

~~~
Scope results: 679
( ( TITLE-ABS-KEY ( music ) OR TITLE-ABS-
KEY ( music therapy ) ) AND ( TITLE-ABS-KEY ( chronic pain ) OR TITLE-ABS-KEY ( neuropathic pain ) OR TITLE-ABS-
KEY ( peripheral neuropathy ) OR TITLE-ABS-KEY ( neuralgia ) OR TITLE-
ABS-KEY ( central pain ) OR TITLE-ABS-KEY ( back pain ) OR TITLE-ABS-
KEY ( irritable bowel ) OR TITLE-ABS-
KEY ( inflammatory bowel ) OR TITLE-ABS-
KEY ( chronic pelvic pain ) OR TITLE-ABS-
KEY ( cardiovascular pain ) OR TITLE-ABS-KEY ( cancer ) OR ( TITLE-
ABS-KEY ( tumor ) AND TITLE-ABS-KEY ( pain ) ) OR ( TITLE-ABS-
KEY ( tumour ) AND TITLE-ABS-KEY ( pain ) ) OR TITLE-ABS-
KEY ( phantom limb ) OR TITLE-ABS-
KEY ( complex regional pain ) OR TITLE-ABS-KEY ( *arthritis ) OR TITLE-ABS-KEY ( systemic lupus erythematosus ) OR TITLE-ABS-
KEY ( multiple sclerosis ) OR TITLE-ABS-KEY ( fibromyalgia ) OR TITLE-ABS-KEY ( ankylosing spondylitis ) OR TITLE-ABS-
KEY ( headache ) OR TITLE-ABS-KEY ( palliative care ) OR TITLE-ABS-KEY ( carpal tunnel syndrome ) OR TITLE-ABS-
KEY ( glaucoma ) ) ) AND ( ALL ( randomized controlled trial ) ALL ( randomi sed controlled trial ) OR ALL ( rct ) OR ALL ( controlled clinical trial ) OR ALL ( random allocation ) OR ALL ( controlled trial ) ) OR ALL ( clinical trial )
~~~

For PsycINFO:

(music OR music therapy) AND (chronic pain OR neuropathic pain OR peripheral neuropathy OR neuralgia OR central pain OR back pain OR irritable bowel OR inflammatory bowel OR chronic pelvic pain OR cardiovascular pain OR cancer OR tumor OR tumour OR phantom limb OR complex regional pain OR arthritis OR systemic lupus erythematous OR multiple sclerosis OR fibromyalgia OR ankylosing spondylitis OR headache OR palliative care OR carpal tunnel syndrome OR glaucoma) AND (randomised controlled trial OR rct OR controlled clinical trial OR random allocation OR controlled trial OR clinical trial)

